# Senescent response in inner annulus fibrosus cells in response to TNFα, H_2_O_2_, and TNFα-induced nucleus pulposus senescent secretome

**DOI:** 10.1101/2022.12.21.521533

**Authors:** Aaryn Montgomery-Song, Sajjad Ashraf, Paul Santerre, Rita Kandel

**Author notes:** Corresponding author (RK).

## Abstract

Senescence, particularly in the nucleus pulposus (NP) cells, has been implicated in the pathogenesis of disc degeneration, however, the mechanism(s) of annulus fibrosus (AF) cell senescence is still not well understood. Both TNFα and H_2_O_2_, have been implicated as contributors to the senescence pathways, and their levels are increased in degenerated discs when compared to healthy discs. Thus the objective of this study is to identify factor(s) that induces inner AF (iAF) cell senescence. Under TNFα exposure, at a concentration that can induce senescence in NP cells, bovine iAF cells did not undergo senescence, indicated by their ability to continue to proliferate as demonstrated by Ki67 staining and growth curves and lack of expression of the senescent markers, p16 and p21. Unlike iAF cells, NP cells treated with TNFα accumulated more intracellular ROS and secreted more H_2_O_2_. Following TNFα treatment, only iAF cells had increased expression of the superoxide scavengers *SOD1* and *SOD2* whereas NP cells had increased *NOX4* gene expression, an enzyme that can generate H_2_O_2_. Treating iAF cells with low dose H_2_O_2_ (50 μM) induced senescence, however unlike TNFα, H_2_O_2_ did not induce degenerative-like changes as there was no difference in *COL2, ACAN, MMP13*, or *IL6* gene expression or number of COL2 and ACAN immunopositive cells compared to untreated controls. The latter result suggests that iAF cells have distinct degenerative and senescent phenotypes. To evaluate paracrine signalling, iAF and TNFα-treated NP cells were co-cultured. In contact co-culture the NP cells did induce iAF senescence. Thus, senescent NP cells may secrete soluble factors that induce degenerative and senescent changes within the iAF. This may contribute to a positive feedback loop of disc degeneration. It is possible these factors may include H_2_O_2_ and cytokines (TNFα). Further studies will investigate if human disc cells respond similarly.

## Introduction

Intervertebral disc (IVD) degeneration has a lifetime prevalence of up to 80% and is associated with the development of back pain(1), one of the most common causes of disability in Canada leading to millions of dollars in health care costs and lost wages(2,3). Despite the prevalence of IVD degeneration, the etiology and pathogenesis of degeneration is still poorly understood. More recently, an association between the pathological changes in the IVD and the presence of senescent cells has been identified(4–6). Previous studies have found significantly more senescent cells in human disc tissue herniations(4) than healthy discs. Similarly, aged or degenerative mouse(7) and rat(8) IVDs accumulate significantly more senescent cells than healthy discs. Interestingly, a p16 knockout mouse, a protein identified as a key driver of the senescence program leads to amelioration of specific markers of IVD degeneration(7,9). Further, treatment with senolytic drugs have been shown to reduce IVD degeneration severity(10,11). This has led many to believe that cellular senescence plays a role in the pathophysiology of IVD degeneration.

The intervertebral disc is composed of a nucleus pulposus surrounded by annulus fibrosus which can be divided further into an inner and outer zone based on the composition. The inner annulus is integrated with the nucleus pulposus. There is a high degree of variability in the reported senescence rate within the tissues of the IVD in humans, ranging from 13-86% in the NP(4,8,9,12) and 5-86% in the AF(8,13,14). This variation likely reflects the method of senescence identification. One of the most well studied inducers of senescence in NP cells is TNFα(15), however the mechanism through which TNFα induces senescence is still not fully delineated. Studies have implicated PI3K-Akt(15), and pSTAT3(16) as potential signaling mechanisms. Recent work has demonstrated that TNFα induced senescent NP cells secrete soluble factors that are capable of inducing senescence in healthy NP cells(16), however the effect of these soluble factors on AF cells have not been assessed. A further understanding of the impact of the NP senescent secretome on other disc cells is important to enable understanding of how the propagation of the degenerative phenotype within the disc occurs.

Reactive oxygen species (ROS), i.e., superoxide and H_2_O_2,_ have also been shown to be capable of inducing senescence in a number of different cell types, including NP cells(17) and AF cells(18). H_2_O_2_ is a redox signaling factor and is produced by normal metabolizing cells. The H_2_O_2_ concentration in the cell is in the nanomolar concentration and outside of the cell ranges from approximately 1-5 μM(19). It signals, in part, by reversible oxidation of specific protein Cys thiolate residues which activates redox signalling. These can then activate phosphorylation cascades and transcription, to name a few processes(20). H_2_O_2_ is produced through multiple pathways, one of which is the NADPH oxidase (NOX) family and the complexes of the electron transport chain(21). NOX2 and 4 expression has been shown to increase within the IVD during degeneration in rats(22), and in NP cells following exposure to IL1α or ROS in-vitro(23,24). Mitochondrial dysfunction has also been associated with IVD degeneration and has been proposed to contribute to ROS-induced damage within the tissues of the IVD(25). In response to oxidative stress, mammalian cells have four enzymes that compose the primary ROS response: superoxide dismutase (SOD), catalase (CAT), glutathione peroxidase (GPX)(26), and peroxiredoxins (PRX). SOD catalyzes the dismutation of superoxide radicals to H_2_O_2_ which can then be degraded into H_2_O via CAT, GPX, or PRX. Cells in the intervertebral disc have been reported to express all isoforms of SOD(27,28), although studies have consistently found a higher expression of SOD in the AF when compared to the NP(28,29). Decreased SOD and catalase activity/expression has been associated with IVD pathologies(30), which has led many to believe that a redox imbalance within the IVD may contribute to the pathogenesis of IVD degeneration(31). Despite this, characterization of ROS scavenging systems within the disc remains poorly understood and has not been determined in the iAF.

Thus, the objective of this study is to identify factor(s) that induces iAF cell senescence. These studies will provide insight into factor(s) leading to iAF senescence and the contribution of inter-tissue communication on this process.

## Methods

### Cell isolation and monolayer culture

Intervertebral discs were aseptically excised from bovine caudal spines. IAF and NP tissues were visually distinguished and harvested as previously described(32–34). NP and iAF tissues were each finely diced in HAM’s F12 (Wisent, 318-010-CL) into approximately 5 mm^3^ cubes. Tissues were individually digested in 0.3% protease (Type XIV, P5147, Sigma-Aldrich, St Louis, MO, USA) in HAM’s F12 supplemented with 100 U/mL of penicillin-streptomycin (Gibco, 15140122) at 37°C for 1 hour, followed by 0.2% collagenase A (COLLA-RO, Sigma-Aldrich, St Louis, MO, USA) in HAM’s F12 supplemented with 100 U/mL of penicillin-streptomycin (Gibco, 15140122) at 37°C for approximately 16 hours. Digested tissues were passed through 100 μm cell strainer and centrifuged at 800g for 8 minutes. Cells were washed three times in DMEM (Wisent, 319-016-CL) supplemented with 5% fetal bovine serum (Wisent Bioproducts, St-Bruno, Quebec, Canada). Approximately 17,000 cells/cm^2^ of primary (P0) NP or iAF cells were plated separately in monolayer culture in T175 flasks (Sarstedt, 83.3912.502) in DMEM supplemented with 5% fetal bovine serum (FBS). P0 iAF and NP cells were cultured for 6 days and then passaged. Passage 1 cells were used in all experiments, unless otherwise stated.

In selected experiments, cells were treated with TNFα (40 ng/mL, R&D systems recombinant bovine TNFα 2279-BT, reconstituted in 0.1% bovine serum albumin in PBS) in DMEM supplemented with 5% FBS for 24 hours, followed by 5 washes with DMEM and then cultured cytokine-free in DMEM containing 5% FBS and allowed 24 hours to recover, prior to analysis.

### Non-contact co-culture

IAF and NP cells were isolated and cultured as described above. Monolayer P1 iAF and NP cells were used for all non-contact coculture experiments. Approximately 14,000 cells/cm^2^ iAF cells were seeded onto 12-well plates (Sarstedt, 83.3921.005) and in separate cultures approximately 14,000 cells/cm^2^ NP cells were seeded onto hanging inserts (Corning Costar 0.2 µm pore PTFE transwells). This cell density was used so that cells were not confluent at the time of analysis. NP cells were treated with 40 ng/mL of TNFα for 24 hours, followed by 5 washes with DMEM only. Transwell inserts were then placed in the wells containing iAF cells and cocultured for 24 hours in DMEM containing 5% FBS prior to analysis.

### Contact Co-Culture

Contact coculture was performed as previously described(16) with some modifications. DMEM containing 5% FBS was used for all contact cocultures, unless otherwise stated. NP cells were passaged to P1, plated at 14,000 cells/cm^2^ and treated with TNFα (40 ng/mL) for 24 hours. The iAF cells (P0) were cultured for 5-7 days until harvested using trypsin-EDTA for experimental set up. NP cells (P1) were trypsinized and resuspended in 20 μM CellTracker Red CMTPX dye (ThermoFisher, C34552) in DMEM (Wisent, 319-016-CL) according to the manufacturer’s instructions. Greater than 95% labelling of cells was confirmed by fluorescent microscopy. NP and iAF cells were then mixed at a ratio of 1:1 and plated into chamber slides at approximately 14,000 cells/cm^2^ (7,000 cells/cm^2^ of each iAF and NP, or 14,000 cells/cm^2^ of iAF alone) (Ibidi, 81816). Experimental conditions were as follows: iAF cells alone, untreated NP and iAF coculture, and TNFα treated-NP and iAF coculture. In the final cocultures, NP cells were P2, and iAF cells were P1. Cells were cultured for 24 hours prior to analysis. Cells were evaluated for senescence by p16 immunocytochemistry as described below (Roche, CINtec 06695248001). Cells were imaged using a Leica DMI-6000 spinning disc confocal microscope running Velocity imaging software. All contact coculture quantification was assessed manually. P16 immunostaining was visualized in the far-red channel, NP cells were labelled with CellTracker Red/DAPI, and iAF cells were positive for DAPI but were negative for CellTracker Red. Each image was assessed for number of p16^+^ iAF, p16^+^ NP, total iAF, and total NP cells. A minimum of 100 cells were assessed for each biological replicate. Results were displayed as the percentage of p16 positive iAF and NP cells in each condition.

### Formation of 3D tissue sheets

Tissue sheets were formed by seeding P1 iAF cells at high density (570k cells/cm^2^) in 12-well plates (Sarstedt, 83.3921.005). Tissues were cultured in DMEM supplemented with L-Proline (40 μg/mL, Sigma-Aldrich, St. Louis, MO, USA), Insulin-Transferrin-Selenium (1%, Wisent Bioproducts, St-Bruno, Quebec, Canada), sodium pyruvate (1 mM, Wisent Bioproducts, St-Bruno, Quebec, Canada), and 10% fetal bovine serum (complete medium). Non-adherent cells were removed after 2 days and replaced with complete medium with ascorbic acid (100 μg/mL, Sigma-Aldrich, St. Louis, MO, USA). The media was replaced with fresh complete media with ascorbic acid every other day. Tissues were harvested after 10 days of culture. In selected experiments the iAF tissue sheets on day 9 of culture were treated for 24 hours with TNFα (40 ng/mL) or 50 μM H_2_O_2_, followed by 5 washes with DMEM. The cultures were then placed in complete media without ascorbic acid and harvested 24 hours later.

### Histology and immunofluorescence

Monolayer cells were fixed in 4% paraformaldehyde (PFA) for 10 minutes and tissue sheets were fixed in 10% formalin for 12 minutes for histology and immunofluorescence and placed in 30% sucrose. Using a dissection microscope, agarose covered cell sheets were cut into thirds (each 12 mm wide). The center third of the tissue was mounted in OCT and cut cross-sectionally at 7 μm using a cryostat. Sections were collected onto silane coated slides and dried overnight at 40°C.

Sections were stained with hematoxylin and eosin, or Toluidine blue and cover-slipped using Micromount (Leica Biosystems, Buffalo Grove, IL USA). Tissues were imaged with a light microscope (Olympus BX61) running CellSens version 1.18.

Collagen type 1 and 2 immunohistochemistry was preceded by antigen retrieval using enzymatic digestion. Sections were incubated in Tris-buffered saline (TBS, pH 2) for 5 minutes, followed by pepsin (2.5 mg/mL in TBS pH2, Millipore Sigma P7012) for 10 minutes at room temperature, followed by 3 washes with PBS. For aggrecan immunostaining the sections were incubated in hyaluronidase (25 mg/mL in PBS pH 7.3, Millipore Sigma H3506) for 30 minutes at 37°C. Boiling sections in Dako Target Retrieval solution, pH 9.0 (Agilent, S236784-2) for 10 minutes was used for MMP13 and p16 antigen retrieval. Monolayer cell staining did not utilize any antigen retrieval.

Sections or cells were then washed three times with PBS and blocked in 20% goat serum (Gibco, 16210-064) and 0.1% Triton X-100 in PBS. Sections were incubated with primary antibody (listed in S3 methods) at 4°C overnight in a humidified chamber. The sections were washed three times with PBS before incubating with AlexaFluor secondary antibody (listed in S3 methods) and 4’6-diamidino-2-phenylindole (DAPI) together at room temperature for 1 hour. Sections were washed 5 times with PBS, stained with DAPI and mounted with PermaFluor™ (Thermofisher Scientific, Waltham, MA, USA). Negative controls consisted of replacing the primary antibody with a species matched IgG antibody at the same protein concentration (w/v). Immunofluorescence was imaged with a fluorescent microscope (Olympus IX81) and Velocity version 6.3.0. Quantification of ECM proteins in monolayer were assessed using ImageJ version 1.53q. COL1, COL2, p16, and ACAN were all stained independently and viewed in the red channel. The images were captured from the same 3 locations in each well using a well-overlay method (S3 methods). Nuclear counting was automated by converting the image to binary and watershed separation. Positive cells (red cytoplasm) were counted manually.

Quantification of tissue thickness was calculated by measuring the average distance between the upper and lower cross-sectional edges at 4 standard sites in the tissue in ImageJ version 1.53q. 2 sections approximately 0.2 mm apart and 2 images per section were assessed for all tissue analysis. 3 biological and 2 technical replicates were used for all tissue sheet quantification assays.

### Growth curves

To assess monolayer cell proliferation growth curves were determined over 72 hours, iAF cells were seeded in chamber slides at approximately 14,000 cells/cm^2^(Ibidi, 81816). Following 24 hours of treatment with TNFα or serum starvation, iAF cells were fixed at 24 hour intervals over 3 days with 4% PFA. Cells were stained with DAPI for 15 minutes in PBS. All cells were imaged in each well with a fluorescent microscope (Olympus IX81) and Velocity version 6.3.0. Nuclear counting was automated using ImageJ by converting the image to binary and watershed separation. All the cells were counted within the wells of each replicate.

### Senescence associated beta-galactosidase staining

Senescence associated beta-galactosidase activity (SA-βGal) was evaluated using the SA-βGal staining kit (#9860, Cell Signaling Technology, Danvers, MA USA) according to the manufacturer’s directions. Briefly, cells or tissues were fixed using solution composed of 2% formaldehyde/0.2% glutaraldehyde in PBS for 15 minutes. Cells were washed 3 times in PBS and incubated with the staining solution adjusted to pH 6 (40 mM citric acid/phosphate buffer, 5mM K_4_[Fe(CN)_6_] 3H_2_O, 5mM K_3_[Fe(CN)_6_], 150 mM sodium chloride, 2 mM magnesium chloride and 1 mg ml^−1^ X-gal) in distilled water for 16 hours at 37°C in a non-humidified oven. Cells were washed once with PBS, mounted with 70% glycerol and imaged under phase contrast microscopy (Olympus BX61 microscope running CellSens version 1.18. 3). The images were captured from the same location in each well using a well-overlay method (S3 methods). Any blue stained (SA-β-galactosidase-positive) cells were considered positive. ImageJ was used to count cells and the percentage of SA-β-galactosidase-positive cells (blue stained) was calculated.

### Quantification of secreted H_2_O_2_ by AmplexRed

Cells were seeded in 96-well plates at approximately 14,000 cells/cm^2^. Cells were cultured for 24 hours. In select experiments, cells were pretreated with the NOX inhibitor, 5 μM diphenyleneiodonium chloride (DPI) (Millipore-Sigma, D2926, resuspended to 5 mM in DMSO) in Hanks Balanced Salt Solution (HBSS) for 3 hours. Cells were subsequently treated with 50 μM H_2_O_2_ or 40 ng/mL TNFα for 16 hours. Cells were washed 3 times with DMEM and cultured treatment-free for 24 hours in DMEM containing 5% FBS. The media was removed and 105 μL of PBS was placed on the cells and placed back in the incubator for 1 hour (37C; 5% CO_2_). The PBS supernatant was collected and H_2_O_2_ was quantified using AmplexRed assay (ThermoFisher, A22188) according to the manufacturer’s instructions. An H_2_O_2_ standard curve was created (0.0156 to 2 μM in PBS). Solutions were incubated with AmplexRed solution for 30 minutes in a black, flat bottom, 96-well plate (Caplugs/Evergreen, 290-895-Z1F) at room temperature in the dark, and fluorescence intensity measured using EnSpire 2300 Multilabel Reader (running EnSpire Manager version 2.00) at 560/590 nm.

### CellROX green and JC-1 staining

To assess intracellular ROS and mitochondrial membrane potential CellROX green (ThermoFisher, C10444) and JC-1 (ThermoFisher, T3168) molecular probes were used, respectively. P1 NP and iAF cells were plated at approximately 17,000 cells/cm^2^ (6k cells/well) in 18-well Ibidi chambers (Ibidi, 81816). Cells were treated with TNFα (40 ng/mL) or H_2_O_2_ (50 μM) for 16 hours, then washed three times with DMEM and cultured for 24 hours in DMEM supplemented with 5% FBS. Cells were then incubated with either CellROX green (5 μM) or JC-1 (5 μg/mL) according to the manufacturer’s directions for 1 hour and visualized by epifluorescent microscopy (Olympus BX61). Quantification of CellROX green and JC-1 was done using ImageJ. CellROX green was analyzed for total fluorescence divided by the total number of cells. JC-1 was analyzed for average red fluorescence divided by average green fluorescence.

### Live/Dead and TUNEL staining

To assess viability of iAF or NP cells following exposure to TNFα, H_2_O_2_, and DPI, P1 NP and iAF cells were plated at approximately 14,000 cells/cm^2^ (6k cells/well) in 18-well Ibidi chambers (Ibidi, 81816). Cells were pre-treated with DPI (5μM) or M40403 (100 μM) (Cayman Chemicals, 10 mM in ethanol) resuspended in HBSS for 1 hour, washed 3 times with DMEM, followed by TNFα (40 ng/mL) or H_2_O_2_ (50 μM) in DMEM for 16 hours. In experiments using DPI or M40403, control cells were pre-treated with the same amount of carrier (HBSS).

For the Live/Dead assay (Invitrogen, L3224), cells were washed in DMEM and treated with Calcein-AM (2 μM) and Ethidium homodimer-1 (2 μM) diluted in DMEM for 30 minutes and visualized by epifluorescent microscopy (Olympus BX61). 50 mM H_2_O_2_ diluted in DMEM was used as a positive control.

For the TUNEL apoptosis assay (Terminal deoxynucleotidyl transferase (TdT) dUTP Nick-End Labeling (TUNEL) assay; Roche, 1168479591), cells in monolayer culture were stained according to the manufacturer’s instructions. Cells were washed with PBS once and fixed with 2% PFA for 1 hour. Cells were rinsed once with PBS and permeabilized with the included permeabilization solution for 2 minutes on ice. Label solution and enzyme solution were combined and incubated on cells for 1 hour at 37°C. Cells were rinsed three times with PBS and mounted with PermaFluor™ (Thermofisher Scientific). Cells were incubated with the nuclear stain DRAQ5 (10 µM diluted in PBS, Thermofisher Scientific, 62251) for 15 minutes and visualized by epifluorescent microscopy (Olympus BX61).

### Gene expression

Cells in monolayer were placed in TRIzol (ThermoFisher, 15596026) and RNA isolated according to the manufacturer’s instructions. For the 3D cultures, tissues were rinsed once with PBS and collected into TRIzol (1 mL, 12 well plate), vortexed briefly, and incubated for 10 minutes at which point the tissue was completely dissolved. RNA was isolated according to manufacturer’s instructions. The pellet was washed with 75% ethanol overnight at -20°C. The following day, samples were spun at 7,500g for 5 minutes at 4°C. A second wash was performed with 75% ethanol and samples were air dried for 15 minutes and resuspended in 20 µL nuclease free water. RNA quantity and quality was assessed by spectrophotometer. Reverse transcription was performed using SuperScript III reverse transcriptase (ThermoFisher, 18080093) and 2 µg of RNA, according to the manufacturer’s instructions. qPCR was performed using a Roche LightCycler 96. Primers are listed in S3 methods. Gene expression analysis was presented as 2^-ΔCt^. To calculate ΔCt, technical replicates were averaged, and average 18S rRNA Ct values were subtracted from average Ct values of the gene of interest from the same biological replicate.

### Statistics

At least 3 biological replicates were obtained for each experiment, and 3 technical replicates/condition were performed unless otherwise specified. One biological replicate was composed of tissue from 3 intervertebral discs of a single bovine caudal spine. Unpaired T-test was used when comparing between two groups, and one-way or two-way analysis of variance (ANOVA) was used when comparing multiple conditions. To minimize family-wise type I error, Tukey’s HSD post-hoc test was used when comparing multiple means. Significance was defined as p<0.05. Analysis was done using GraphPad Prism Version 9.2.0.

### Ethics Statement

As the tissue was obtained from the abbatoir after euthanasia and is considered waste, no REB was required.

## Results

### TNFα induces an altered phenotype but not senescence in iAF cells

TNFα treated iAF cells had significantly more senescence associated β-galactosidase (SA-βGal) positive cells as compared to control (Fig 1A). However, it did not induce senescence as they retained their ability to proliferate as indicated by quantifying cell number and Ki67 immunostaining over a 3-day period (Fig 1B-D, S1 Fig). TNFα exposure did not increase p21 or p16 accumulation, as determined by immunostaining (Fig 1E/F).

**Fig 1:**
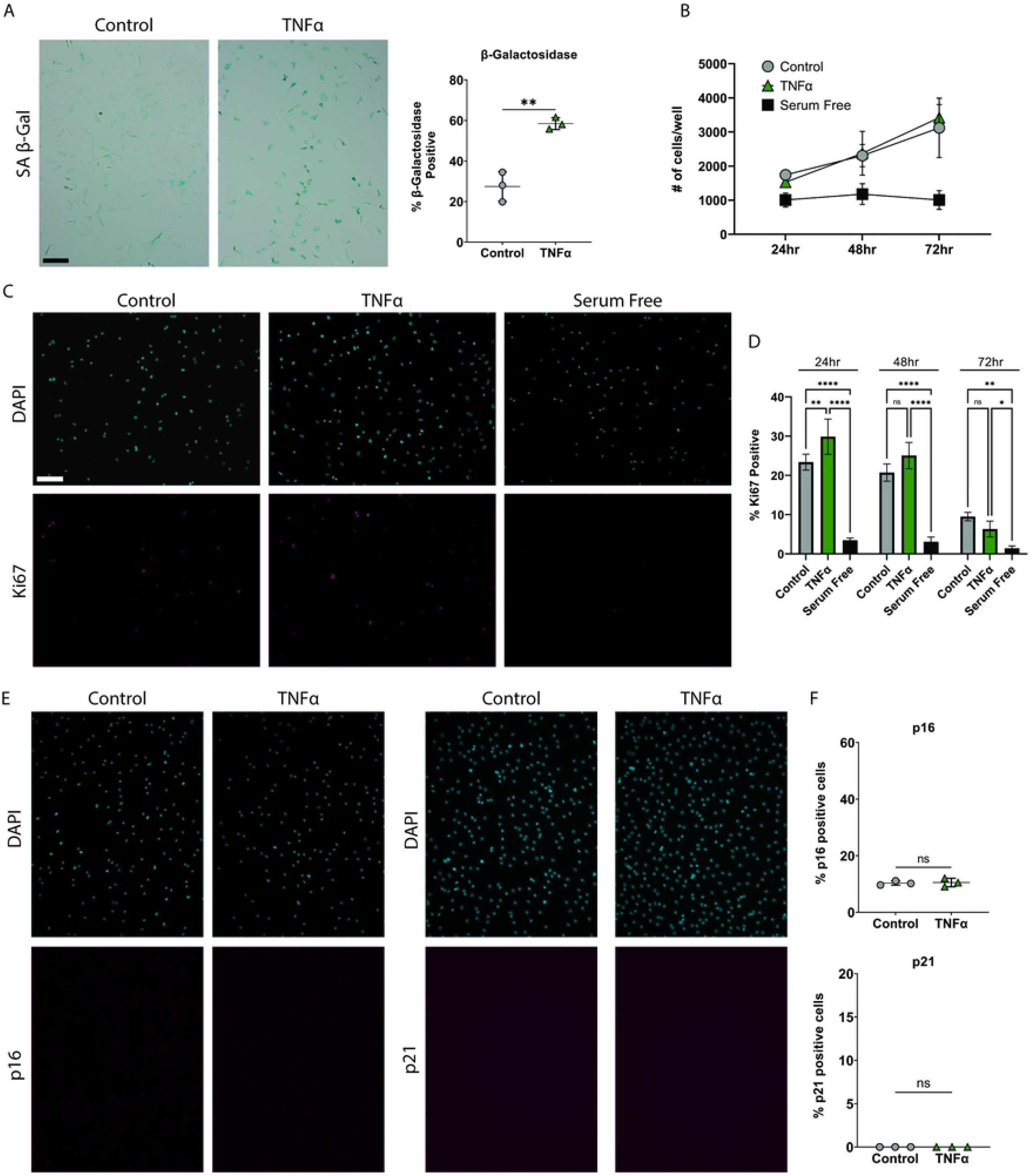
iAF cells are resistant to TNFα induced senescence. **(A)** Representative phase contrast images and quantification of senescence associated β-galactosidase staining of iAF cells treated with TNFα. **(B)** Growth curve of iAF cells treated with TNFα or serum starved at 24, 48, and 72 hours. **(C)** Representative images of Ki67 immunocytochemistry. **(D)** Quantification of percentage of Ki67 positive cells at 24, 48, and 72 hours post treatment. Media was not changed throughout the 72 hour time course. **(E)** Representative images of p16 and p21 immunostaining of iAF cells treated with TNFα. **(F)** Quantification of p16 and p21 immunostaining of iAF cells treated with TNFα, represented as percentage of total cells that were stained. Scale bar = 100μm. p <0.05 = *, p<0.01 = **, p<0.001 = ***, p<0.0001 = ****, N=3 for immunostaining, N=4 for gene expression.

### iAF and NP cells have a differential ROS response following exposure to TNFα

TNFα induced a change in ROS response in NP cells, with an increase in intracellular ROS as demonstrated by CellROX green staining (Fig 2A/B) and H_2_O_2_ secretion as quantified by AmplexRed assay (Fig 2C). However, unlike NP cells, iAF cells exposed to TNFα showed no increase in intracellular ROS accumulation or H_2_O_2_ secretion (Fig 2D-F).

**Fig 1:**
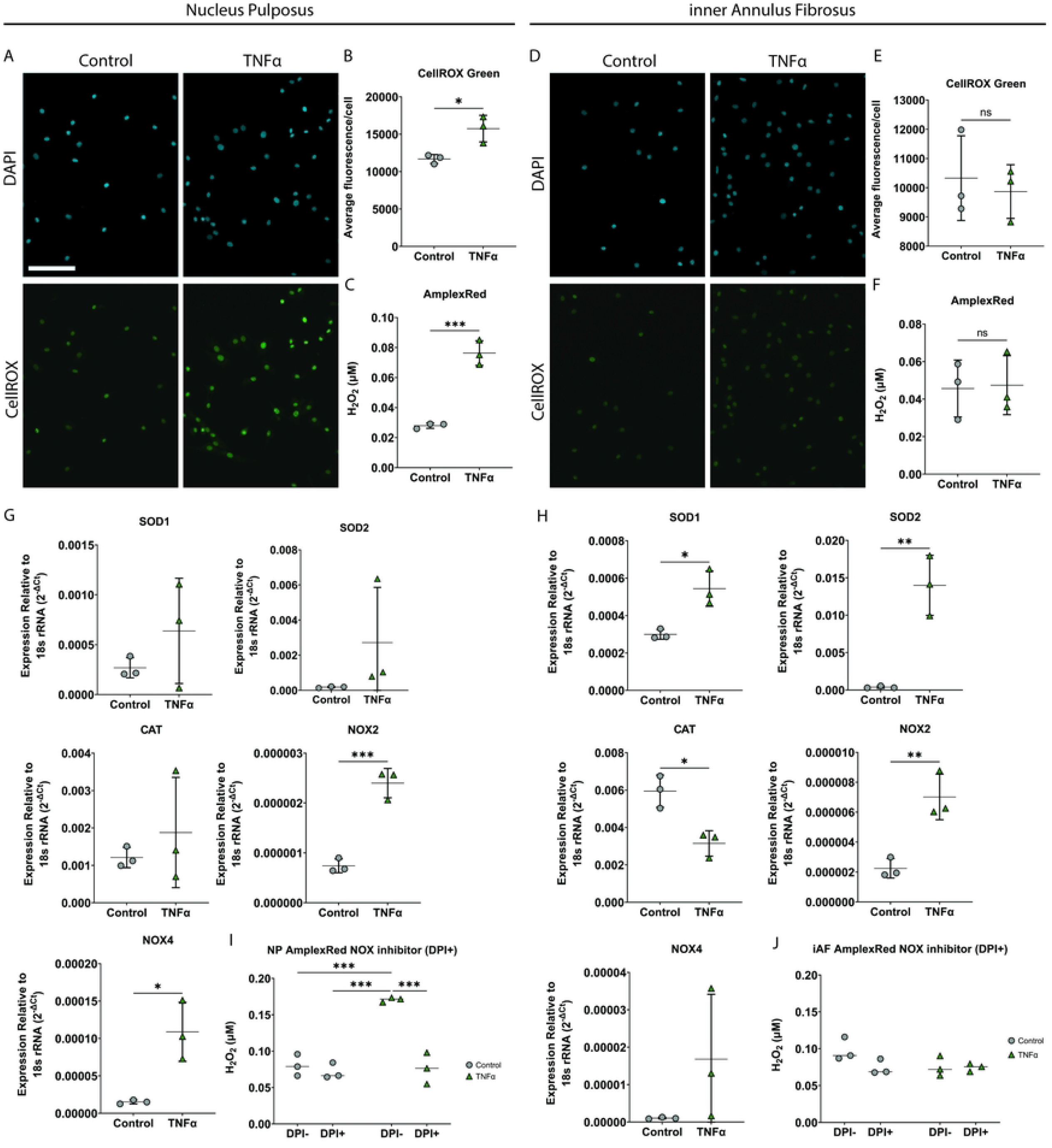
iAF cells have a differential ROS response to TNFα compared with NP cells. **(A)**Representative images of CellROX green staining in NP cells exposed to TNFα (40 ng/mL). **(B)** Quantification of CellROX green average fluorescence intensity per cell in NP cells exposed to TNFα. **(C)** AmplexRed assay quantification of H_2_O_2_ released from NP cells exposed to TNFα. **(D)** Representative images of CellROX green staining in iAF cells exposed to TNFα (40 ng/mL). **(E)** Quantification of CellROX green average fluorescence intensity per cell in iAF cells exposed to TNFα. **(F)** AmplexRed quantification of H_2_O_2_ released from iAF cells exposed to TNFα. **(G)** Gene expression analysis relative to 18s rRNA of ROS related genes in NP cells exposed to TNFα. **(H)** Gene expression analysis relative to 18s rRNA of ROS related genes in iAF cells exposed to TNFα. **(I)** AmplexRed quantification of H_2_O_2_ released from NP cells exposed to TNFα pre-treated with the NOX-inhibitor diphenyleneiodonium chloride (DPI). **(J)** AmplexRed quantification of H_2_O_2_ released from iAF cells exposed to TNFα pre-treated with the NOX-inhibitor DPI. Scale bar = 100μm. p<0.05 = *, p<0.01 = **, p<0.001 = ***, p<0.0001 = ****, N=3 for all experiments.

NP cells had no significant change in Superoxide dismutase *(SOD)-1, SOD2*, or catalase (*CAT)* gene expression upon exposure to TNFα. However, in contrast to the NP cells, iAF cells yielded significantly higher expression of *SOD1* and *SOD2* following TNFα treatment when compared to their respective untreated cells. Unlike *SOD, CAT* expression was higher in untreated iAF cells than in TNFα treated iAF cells. NADPH oxidase (*NOX) 1-5* gene expression was also assessed, however, only *NOX2* and *NOX4* were detectable in both the NP and iAF cells. *NOX2* but not *NOX4* expression was higher in TNFα treated iAF cells as compared to untreated iAF cells. Both *NOX2* and *NOX4* expression was significantly increased in TNFα treated NP cells when compared to untreated NP cells (Fig 2G/H).

Treating NP cells with diphenyleneiodonium chloride (DPI), a broad NOX inhibitor, caused a significant decrease in TNFα-mediated H_2_O_2_ accumulation (Fig 2I). iAF cells treated with DPI had no significant change in H_2_O_2_ accumulation (Fig 2J).

### iAF cells undergo senescence when exposed to low dose H_2_O_2_

To determine if iAF cells undergo senescence in response to H_2_O_2_, which could explain the lack of response to TNFα, iAF cells were exposed to low dose H_2_O_2_. H_2_O_2_ treated iAF cells in monolayer show increased staining for SA-βGal (Fig 3A), as well as an increased number of p16 and p21 immunopositive compared to control cells (Fig 3B/C). H_2_O_2_ exposure did not induce changes in *SOD1, SOD2, CAT, NOX2, or NOX4* gene expression (Fig 3D). Despite this, iAF cells exposed to H_2_O_2_ had a disrupted redox regulation with an increase in intracellular ROS and secreted H_2_O_2_, evaluated by CellROX green and AmplexRed assay, respectively (Fig 3E/F). H_2_O_2_ treated iAF cells also had depolarized mitochondrial membrane potential compared to untreated and TNFα treated cells, as visualized by JC-1 red/green fluorescence intensity (Fig 3G).

**Fig 1:**
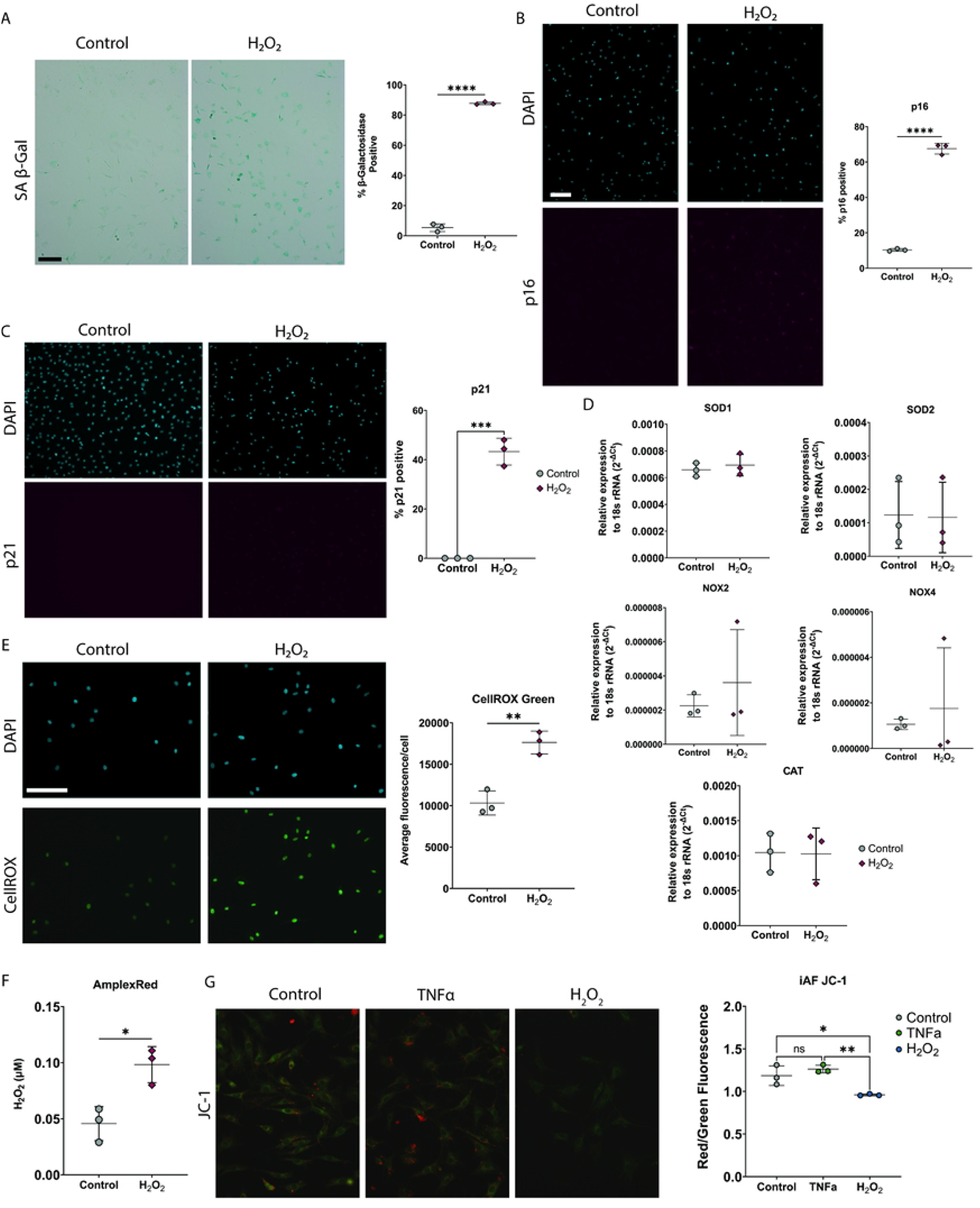
iAF cells are sensitive to H_2_O_2_-induced senescence. **(A)** Representative phase contrast images and quantification of senescence associated β-galactosidase staining of iAF cells treated with H_2_O_2_ (50 μM). **(B)** Representative images and quantification of p16 immunostaining of iAF cells treated with H_2_O_2_. **(C)** Representative images and quantification of p21 immunostaining of iAF cells treated with H_2_O_2_. **(D)** Gene expression analysis of iAF cells treated with H_2_O_2_. **(E)** Representative images and quantification of intracellular ROS with CellROX green. **(F)** AmplexRed quantification of H_2_O_2_ released from iAF cells exposed to H_2_O_2_. **(G)** Representative images and quantification of JC-1 staining in iAF cells exposed to TNFα or H_2_O_2_. Scale bar = 100μm. p <0.05 = *, p<0.01 = **, p<0.001 = ***, p<0.0001 = ****, N=3 for all experiments.

### iAF cells treated with TNFα but not H_2_O_2_ undergo degenerative-like changes

IAF cells in monolayer treated with TNFα (40 ng/mL) showed a significant reduction in *COL2* and *ACAN* and an increase in *IL6* and *MMP13* gene expression (Fig 4A). This correlated with immunohistochemical data showing a significant reduction in the number of cells producing type II collagen and aggrecan when compared to untreated control cells (Fig 4B/C). Unlike TNFα, H_2_O_2_ (50 μM) exposure did not induce changes in *COL1, COL2, ACAN, IL6*, or *MMP13* gene expression (Fig 4D) or COL1, COL2, or ACAN immunopositivity (Fig 4E/F).

**Fig 2:**
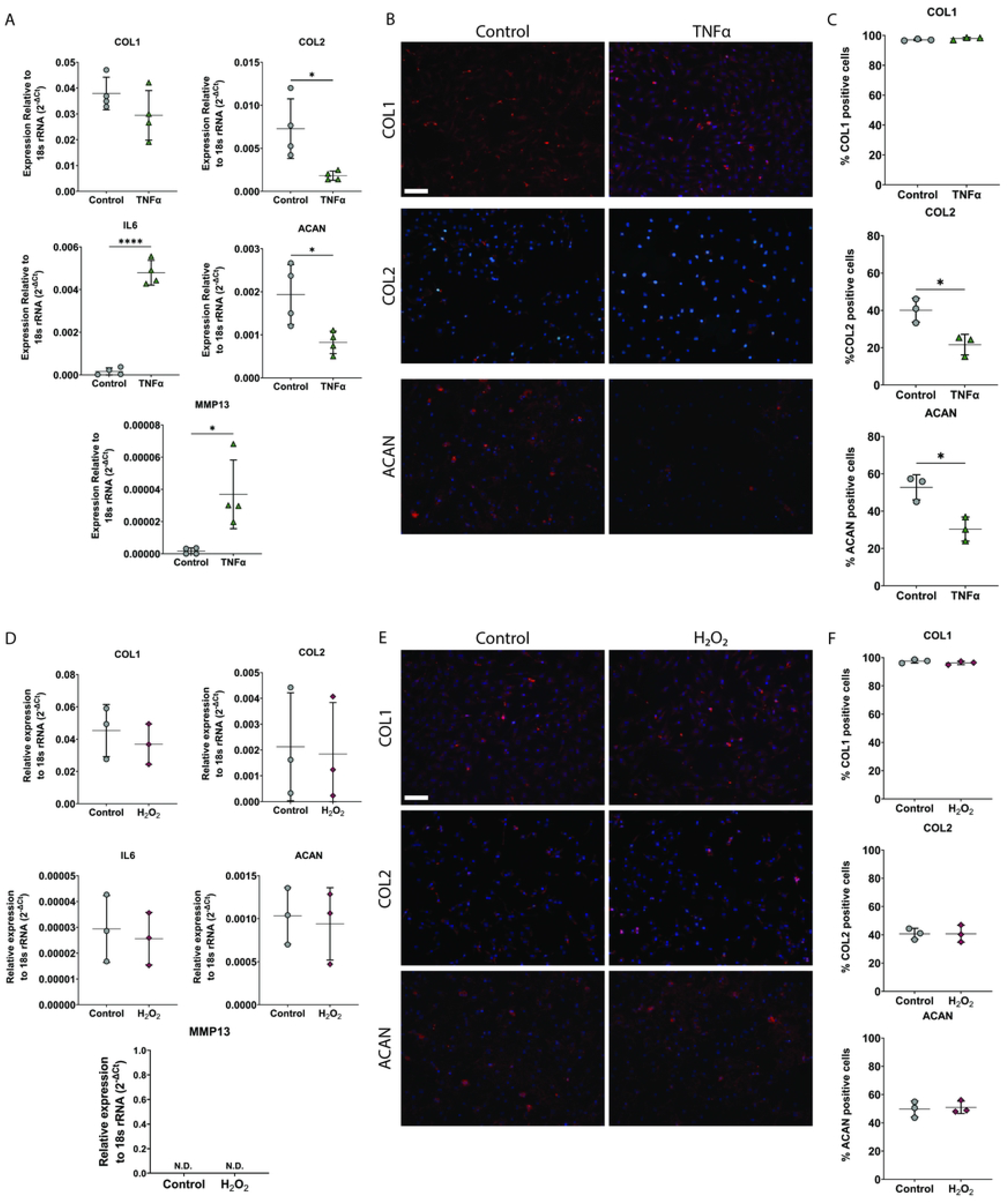
TNFα treated but not H_2_O_2_-induced senescent iAF cells show signs of degeneration at 24hrs. **(A)** Gene expression analysis of iAF cells treated with TNFα (40 ng/mL). **(B)** Representative images of COL1, COL2, and ACAN immunocytochemistry of iAF cells treated with TNFα. **(C)** Quantification of immunocytochemistry in B, presented as percentage of total cells stained. **(D)** Gene expression of iAF cells treated with H_2_O_2_. **(E)** Representative images and quantification of COL1, COL2, and ACAN immunocytochemistry of iAF cells treated with H_2_O_2_. **(F)** Quantification of immunocytochemistry in E, presented as percentage of total cells stained. Scale bar = 100μm. p <0.05 = *, p<0.01 = **, p<0.001 = ***, p<0.0001 = ****, N=3 for all experiments.

### iAF cells grown in 3D tissue sheets also undergo senescence in the presence of H_2_O_2_ but not TNFα

To determine if the inability of TNFα to induce senescence in iAF cells was an artefact of growing the cells in monolayer culture, cells were also grown in 3D to form tissue. These tissues contain collagen types I and II and aggrecan similar to native iAF (Fig 5A). As in monolayer, TNFα treated iAF tissue showed a significant increase in MMP13 protein as determined by immunostaining. The iAF 3D sheets had a significant increase in the number of p16 immunoreactive cells when exposed to H_2_O_2_ but not TNFα (Fig 5B). Although TNFα and H_2_O_2_ treated iAF tissues showed a significant decrease in thickness compared to untreated controls, the decrease was greater in the cytokine treated tissues (Fig 5C).

**Fig 3:**
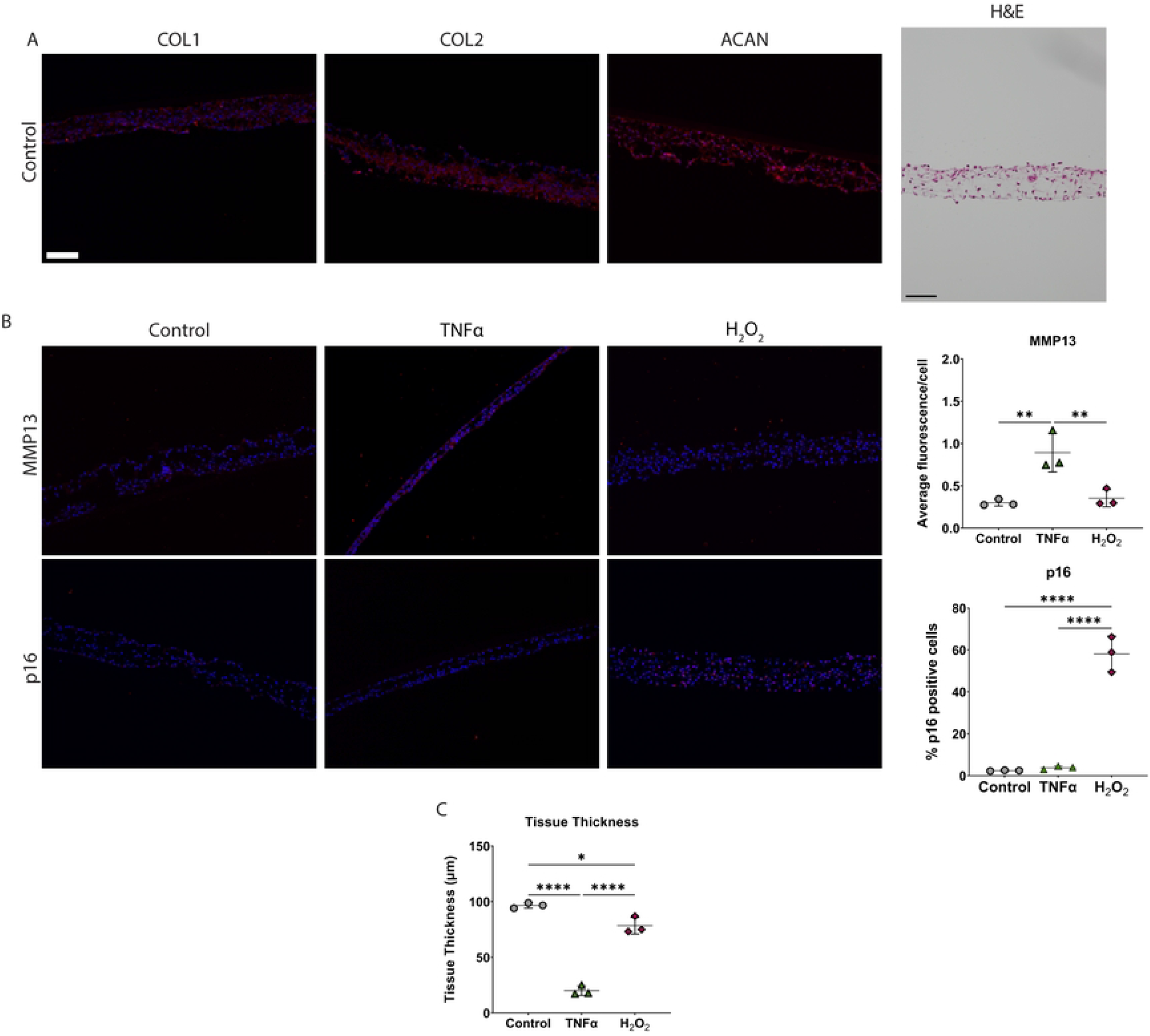
iAF cells in 3D tissue sheets undergo senescence when exposed to H_2_O_2_ but not TNFα. **(A)** Representative images of untreated iAF 3D tissue sheets following immunohistochemical staining for COL1, COL2, or ACAN, or H&E staining. **(B)** Representative images of immunohistochemistry and quantification of MMP13 and p16 of iAF 3D tissue sheets exposed to TNFα (40 ng/mL) or H_2_O_2_ (50 μM) for 24 hours. **(C)** Average cross-sectional thickness of iAF 3D tissue sheets exposed to TNFα or H_2_O_2_. Scale bar = 100μm. p<0.05 = *, p<0.01 = **, p<0.001 = ***, p<0.0001 = ****, N=3.

### iAF cells have an altered phenotype following exposure to media conditioned by TNFα treated-NP cells, but only undergo senescence in a contact co-culture model

To determine if iAF cells can respond to the TNFα treated-NP secretome, iAF cells were co-cultured with either TNFα or untreated-NP cells in a contact or non-contact co-culture system. iAF cells in non-contact co-culture with TNFα treated-NP cells have a significant increase in *IL6* and *MMP13* gene expression (Fig 6A) as well as a reduction in the number of type II collagen immunopositive cells compared to co-culture with untreated-NP cells. (Fig 6B). There was no change in *COL1* or *ACAN* gene and protein expression. IAF cells co-cultured with TNFα-treated-NP also had a significant increase in SA-βGal positive cells (Fig 6C) but were negative for p21 and p16 immunoreactivity (Fig 6D).

**Fig 4:**
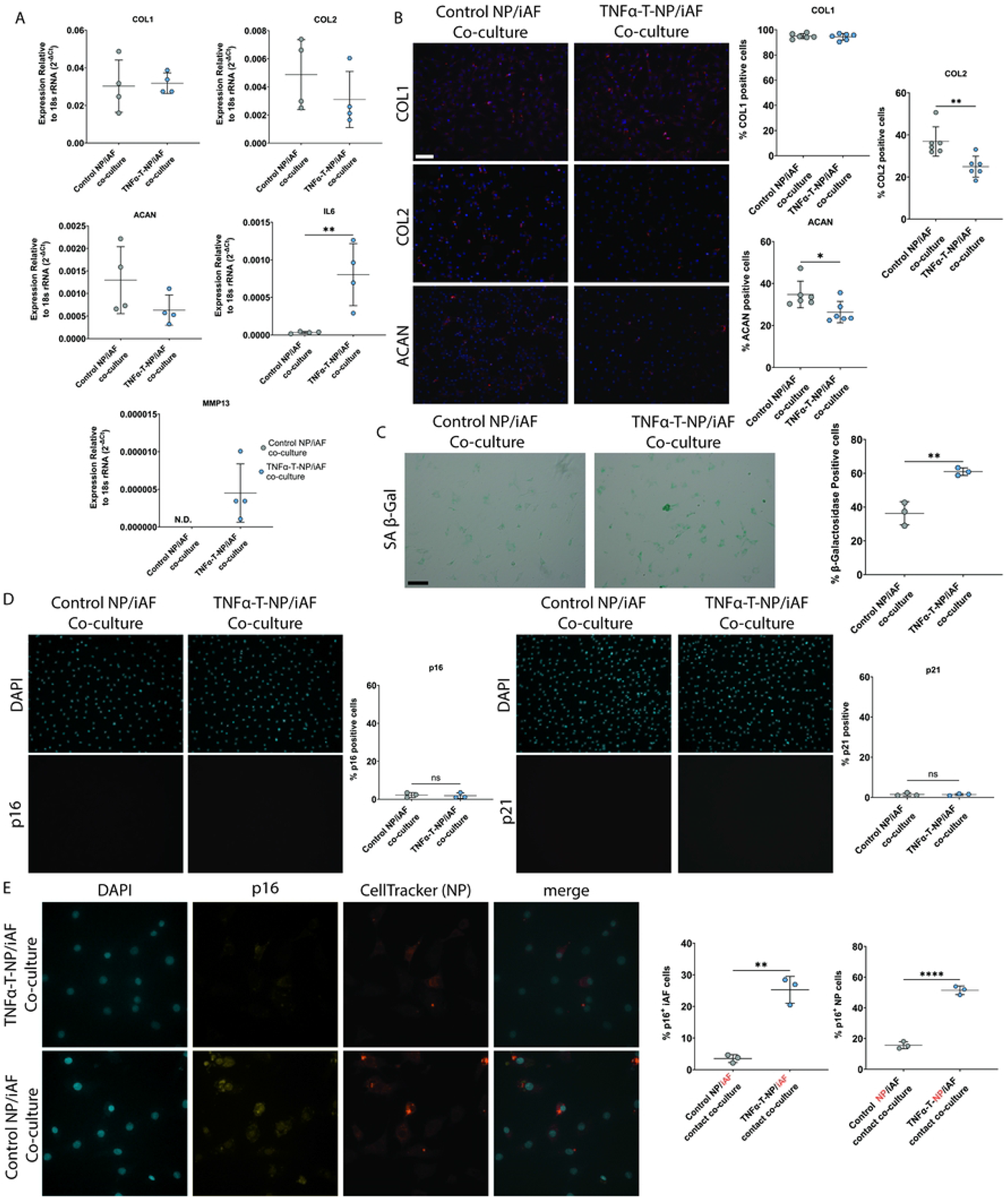
iAF cells have an altered phenotype when co-cultured with senescent NP cells. **(A)** Gene expression of iAF cells exposed to non-contact co-culture with TNFα or untreated NP cells for 24 hours. **(B)** Representative images and quantification of COL1, COL2, and ACAN immunostained iAF cells exposed to non-contact co-culture with NP cells. **(C)** Representative phase contrast images of senescence associated β-galactosidase staining and quantification of iAF cells exposed to non-contact co-culture with NP cells. **(D)** Representative images and quantification of p16 and p12 immunocytochemistry of iAF cells exposed to non-contact co-culture with NP cells. **(E)** Representative images and quantification of iAF cells in a contact co-culture with TNFα and untreated NP cells. Scale bar = 100μm. p<0.05 = *, p<0.01 = **, p<0.001 = ***, p<0.0001 = ****, N=3 for immunostaining, N=4 for gene expression. TNFα-T-NP= TNFα treated NP cells.

In a contact co-culture system, iAF cells admixed with TNFα treated-NP cells, underwent senescence as there was significantly more p16 immunopositive cells when compared to iAF cells cocultured with untreated NP cells (Fig 6E).

## Discussion

Although TNFα is known to induce senescence in NP cells, this study demonstrates that iAF cells are resistant to TNFα-induced senescence under the conditions examined. Previous reports have suggested that increased ROS-accumulation is critical for TNFα-induced senescence(35). Unlike NP cells, iAF cells did not increase intracellular ROS or H_2_O_2_ secretion in response to TNFα. Furthermore, TNFα treated NP cells had increased expression of *NOX4* which has been shown to produce H_2_O_2_(36), whereas TNFα treated iAF cells had increased expression of the superoxide scavengers *SOD1* and *SOD2*. Given that iAF cells were resistant to TNFα Induced senescence, potentially through ROS homeostasis, we next looked to investigate the effect of exogenous low dose H_2_O_2_ on iAF cells. H_2_O_2_ did induce senescence in iAF cells, however unlike TNFα, H_2_O_2_ did not induce the release of markers typically associated with matrix degeneration, as there was no change in *COL2, ACAN, MMP13*, or *IL6* gene expression nor the number of cells producing type II collagen and aggrecan. This suggests that iAF cells may have distinct degenerative and senescent phenotypes. Lastly, as some studies have demonstrated that degenerative changes occur in the NP prior to the AF(37,38) and that senescent NP cells secrete soluble factors capable of inducing senescence in healthy NP cells(16), we next investigated if the senescent NP secretome is capable of inducing senescence in iAF cells. Co-culturing iAF cells with TNFα-induced senescent NP cells, did induce senescence of the iAF cells, as well as inducing some degenerative changes when compared to co-cultures with untreated NP cells. Thus, senescent NP cells may contribute to the senescent and degenerative changes observed within the iAF during IVD degeneration.

Although H_2_O_2_ accumulation within the disc is well characterized(25,39–41), to our knowledge this is the first study to demonstrate that H_2_O_2_, and not the cytokine TNFα, may be a principle driver of senescence in the iAF and importantly, that it may act as a signaling molecule between the NP and iAF. H_2_O_2_ can be transported across the plasma membrane through aquaporins, which are known to be expressed by both NP and AF cells(42,43), and has been proposed to facilitate cell-to-cell signaling(44). The crosstalk between NP and iAF cells is not entirely unexpected as a previous study has demonstrated H_2_O_2_ can be secreted by one cell type and taken up by another in-vitro(44). Given that iAF cells are sensitive to low dose H_2_O_2_, and that TNFα treated NP cells increased the amount of secreted H_2_O_2_, this may suggest that the H_2_O_2_ generated from TNFα-induced senescent NP cells may be responsible for the senescence observed in the co-cultured iAF cells. Interestingly, iAF cells co-cultured with NP cells in a non-contact culture system did not undergo senescence. While there may be the need for direct contact to enable the senescent effects of H_2_O_2_ on iAF cells, it is possible that it just reflects the unstable nature of H_2_O_2_(38). Studies in the literature, using plant cells, demonstrated that H_2_O_2_ diffusion distance within a cell is just on the order of 1μm, with an approximate half life of just 1ms(45,46). If in the non-contact culture H_2_O_2_ levels were not high enough by the time it diffused to the iAF cells or present for sufficient time to induce senescence in the iAF cells under the conditions examined, the effects of H_2_O_2_ may not occur. Alternatively, as H_2_O_2_, or even the ROS generating NADPH oxidase, can be transported via exosomes(47,48), it is possible that diffusion of exosomes was impaired through the transwell pores. Nevertheless, taken together this data indicates that H_2_O_2_ transport within the disc may play a role in senescence propagation between cell types. This contrasts with the dominant theory that propagation of senescence from the NP to AF is that, due to NP degeneration can lead to aberrant ECM remodeling and subsequently compromise of mechanical properties(49). While the latter is likely a contributing cause to the demise of the tissues, since altered biomechanical stimuli within the AF tissues have been shown capable of inducing senescence in AF cells(50), it may not be the initiating factor.

To our knowledge, this is the first report that TNFα may not directly play a role in the induction of iAF senescence. Previous studies have identified resistance to TNFα mediated cytotoxicity, however cellular senescence in those studies was not investigated. It is possible that the mechanism of resistance to TNFα cytotoxicity may be similar to the mechanism of senescence resistance in iAF cells. Resistance to TNFα in embryonic KYM-1 cells appears to be mediated by the loss of TNFα-receptor expression(51) or in human neurons/HeLa cells by secretion of TNFα neutralizing proteins(52). These changes are unlikely occurring in iAF cells, as they still respond to TNFα, as demonstrated by alterations in gene expression and accumulation of ECM components. Alternatively, others have found that differential phospholipase A2 activation altered the ability of TNFα to induce cytotoxicity in epithelial, ovarian, and cervical cell lines(53,54). Phospholipase A2 activation is thought to lead to the production of ROS that can be mitigated by superoxide dismutase (SOD). SOD-mediated protection has been demonstrated in L929 cells and L929.12 cells which are sensitive and resistant to TNFα cytotoxicity, respectively(55). This was validated by studies in epithelial cells(56) where it was shown that SOD2 overexpression protects cells from TNFα-mediated ROS cytotoxicity. Although these studies have not investigated the senescence response of the TNFα-resistant cells, our study has similar findings with respect to the regulation of ROS, potentially via SOD. Further studies are required to fully define the mechanism(s) underlying the senescence resistance of iAF cells to TNFα.

It is not entirely unexpected that NP cells respond differently to TNFα, when compared to iAF cells. Although the NP and iAF cells share some phenotypic similarities, the NP and AF are derived from different embryonic lineages: the NP is from the notochord and the AF from the paraxial mesoderm(57). Furthermore, in adult IVD tissues, these cells still have distinct transcriptomes that may alter their responsiveness to TNFα(29). Previous work has identified SOD2 as one of the top differentially expressed genes between iAF and NP cells, basally and in response to TNFα stimulation(28,29). This difference in *SOD2* is consistent with our findings, as iAF cells were shown to have higher TNFα-induced expression of *SOD1* and *SOD2* as compared to NP cells. The *SOD2* response to TNFα is not found across all cell types, since a reduction in *SOD2* expression in response to TNFα has been reported in mouse embryonic fibroblasts(58). In the current study, evaluation of ROS accumulation from iAF and NP cells showed a significant increase in intracellular ROS in NP but not iAF cells following TNFα -exposure, which is also consistent with iAF cells having greater regulation of superoxide and H_2_O_2_. PG1-FM, an H_2_O_2_ reactive probe(44), also caused higher rates of cell death and had greater fluorescence in the iAF when compared to NP cells (data not shown), which may suggest that iAF cells have higher levels of intracellular H_2_O_2_ – both basally and following TNFα-exposure. Taken together, this suggests that the iAF cells may convert intracellular superoxide to H_2_O_2_ via SODs as a protective mechanism against TNFα induced ROS production, and as the levels are lower subsequently do not induce senescence. There are numerous proteins that regulate cellular ROS effects that were not specifically assessed in this study because of the broad spectrum of their scope. Most notably glutathione-peroxidase, thioredoxin, peroxiredoxin, and glutathione. To date, these molecules have largely been understudied in IVD cells. Specifically, two studies have indicated that AF and NP cells may decrease their expression of GPX and GSH when exposed to ROS stress or in menopause mediated IVD degeneration(59,60). Alternatively, other studies have identified non-coding RNAs such as NKILA/miR-21(61,62), and cellular proteins such as Sirt6(63), or prolonged NF-κB activation can increase resistance to the cytotoxic effects of TNFα(64). This suggests that other mechanisms of TNFα signaling regulation may play a role in the iAF cells resistance to TNFα mediated senescence.

Despite increased senescence associated β-galactosidase positivity in iAF cells following exposure to TNFα or non-contact co-culture NP cells, these cells remained p16 and p21 negative – suggesting they are not senescent. Senescence associated β-galactosidase (SA-βGal) was the first stain reported in the literature to identify senescent cells(65), however on its own it is not sufficient to confirm senescence. The assay is designed to measure the endogenous lysosomal β-galactosidase activity outside the enzyme’s ideal pH range. By using this pH, only cells that express significant amounts of lysosomal βGal are stained. Interestingly, although high SA-βGal activity has been associated with senescence it has been shown that lysosomal βGal is not required for the senescence program(66). Additionally, many studies have found SA-βGal staining in the absence of senescence, such as in high cell density culture and serum starvation(67). Other factors have also been shown to induce a positive βGal stain in the absence of senescence such as tartrate-resistant acid phosphatase (TRAP) expression in osteoclasts(68,69) or lysosomal activity in young neurons(70).

This study has some limitations. Specifically, as this is an in-vitro investigation, the manner by which these cell types interact in this environment may not accurately reflect in-vivo signaling. ROS are capable of reacting with ECM components and integrins(71), which may sequester H_2_O_2_ before it reaches adjacent cell types, although this only enhances the case made for direct cell contact. Similarly, other ligands that may modulate TNFα or ROS responsiveness are known to be sequestered by ECM components, such as TGFβ(72,73). ECM components are also known to alter cellular responsiveness to ligands(74,75) and ROS production(71). Extracellular ROS scavenging enzymes may also inhibit ROS/H_2_O_2_ from acting as a signaling molecule within the disc, which may decrease with increasing degeneration(30). Additionally, the concentrations of TNFα (40ng/mL) and H_2_O_2_ (50µM) used in this study are significantly higher than concentrations found in-vivo (TNFα: 0.05-0.15ng/mL, H_2_O_2_: 0.25μM(12,76)). Finally, NP/iAF cultures were performed at a cell ratio of 1:1, higher than the ratio of NP:iAF in bovine caudal intervertebral discs (NP:iAF ratio in-vivo is approximately 1:2 from cell isolations from bovine caudal spines, data not shown).

In summary, the results suggest that iAF cells are sensitive to the senescent effect of H_2_O_2_. The cells appear to be resistant to TNFα-induced senescence at the concentration evaluated, perhaps due to their inability to produce sufficiently increased H_2_O_2_ levels on exposure to the cytokine. Interestingly, NP cells exposed to TNFα undergo senescence and secrete factors that induce degenerative and senescent changes in iAF cells. Disc degeneration has been shown to be multifactorial, with cytokines, genetics, mechanics, and environmental stressors all contributing to the pathophysiology(77–79). The current study reported here, provides another potential contributing mechanism for the positive feedback loop of disc degeneration, specifically through ROS accumulation and the capacity for NP to signal iAF cells, an action which could promote degenerative and senescent changes. This could inform the choice of IVD degeneration therapeutics. For example, targeting only senescent iAF cells may not fully ameliorate degeneration of this tissue, but reducing NP cell capacity to produce ROS (e.g., NOX knockout), or inflammatory cytokines (e.g., TNFα knockout) to prevent the positive feedback of inflammatory and oxidative stress may be equally if not more efficacious. Further study is required to determine if human disc cells respond similarly.

## Acknowledgements

We would like to thank Drs. Marco Massina and Christopher Chang for providing PG1-FM and insights on ROS signaling. We would also like to thank Ryszard Bielecki and Louise Brown for their microscopy expertise.

## Supplemental Figure Captions

S1 Fig: Representative images of Ki67 immunocytochemistry of iAF cells treated with TNFα (40 ng/mL) or serum starved at 48 and 72 hours of treatment.

S2 Fig: iAF cell viability when exposed to TNFα, H_2_O_2_, and DPI. **(A)** Representative images of TUNEL and ethidium homodimer (red)/Calcein-AM (green) following exposure to TNFα (40 ng/mL) and H_2_O_2_ (50 μM). Positive controls were exposed to DNAse I or for TUNEL staining and 50 mM H_2_O_2_ for ethidium homodimer/calcein-AM staining. **(B)** Quantification of percentage of TUNEL positive iAF cells exposed to TNFα and H_2_O_2_. **(C)** JC-1 staining for mitochondrial membrane potential in NP cells exposed to TNFα and quantification presented as red/green fluorescence intensity. **(D)** Representative images of ethidium homodimer (red)/Calcein-AM (green) stained iAF and NP cells following exposure to diphenyleneiodonium chloride (DPI) (5 μM). Student’s T-test was used for statistical analyses of NP JC-1, one-way ANOVA with Tukey’s post-hoc was used for iAF TUNEL quantification. p<0.05 = *, p<0.01 = **, p<0.001 = ***, p<0.0001 = ****, N=3 for all experiments.

## References

1. Sadowska A, Hausmann ON, Wuertz-Kozak K. Inflammaging in the intervertebral disc. Clinical and Translational Neuroscience [Internet]. 2018 Mar 15 [cited 2022 Jul 24];2(1):1–9. Available from: https://journals.sagepub.com/doi/10.1177/2514183X18761146

2. Rampersaud YR, Power JD, Perruccio A V, Paterson JM, Veillette C, Coyte PC, et al. Healthcare utilization and costs for spinal conditions in Ontario, Canada - opportunities for funding high-value care: a retrospective cohort study. Spine Journal. 2020;20(6):874–81.

3. Martin BI, Deyo RA, Mirza SK, Turner JA, Comstock BA, Hollingworth W, et al. Expenditures and health status among adults with back and neck problems. JAMA - Journal of the American Medical Association. 2008;299(6):656–64.

4. Roberts S, Evans EH, Kletsas D, Jaffray DC, Eisenstein SM. Senescence in human intervertebral discs. European Spine Journal. 2006;15(SUPPL. 3):312–6.

5. Patil P, Niedernhofer LJ, Robbins PD, Lee J, Sowa G, Vo N. Cellular Senescence in Intervertebral Disc Aging and Degeneration. Curr Mol Biol Rep [Internet]. 2018 Dec 25;4(4):180–90. Available from: http://link.springer.com/10.1007/s40610-018-0108-8

6. le Maitre CL, Freemont AJ, Hoyland JA. Accelerated cellular senescence in degenerate intervertebral discs: A possible role in the pathogenesis of intervertebral disc degeneration. Arthritis Res Ther. 2007;9(3):1–12.

7. Patil P, Dong Q, Wang D, Chang J, Wiley C, Demaria M, et al. Systemic clearance of p16INK4a-positive senescent cells mitigates age-associated intervertebral disc degeneration. Aging Cell. 2019;18(3):1–11.

8. Veroutis D, Kouroumalis A, Lagopati N, Polyzou A, Chamilos C, Papadodima S, et al. Evaluation of senescent cells in intervertebral discs by lipofuscin staining. Mech Ageing Dev [Internet]. 2021 Oct 1 [cited 2022 Jul 24];199:1–9. Available from: https://pubmed.ncbi.nlm.nih.gov/34474077/

9. Che H, Li J, Li Y, Ma C, Liu H, Qin J, et al. P16 deficiency attenuates intervertebral disc degeneration by adjusting oxidative stress and nucleus pulposus cell cycle. Elife. 2020;9:80–90.

10. Novais EJ, Tran VA, Johnston SN, Darris KR, Roupas AJ, Sessions GA, et al. Long-term treatment with senolytic drugs Dasatinib and Quercetin ameliorates age-dependent intervertebral disc degeneration in mice. Nat Commun [Internet]. 2021;12(1):1–17. Available from: http://dx.doi.org/10.1038/s41467-021-25453-2

11. Lim S, An SB, Jung M, Joshi HP, Kumar H, Kim C, et al. Local Delivery of Senolytic Drug Inhibits Intervertebral Disc Degeneration and Restores Intervertebral Disc Structure. Adv Healthc Mater. 2022;11(2):1–16.

12. Kim KW, Chung HN, Ha KY, Lee JS, Kim YY. Senescence mechanisms of nucleus pulposus chondrocytes in human intervertebral discs. Spine Journal [Internet]. 2009;9(8):658–66. Available from: http://dx.doi.org/10.1016/j.spinee.2009.04.018

13. Gruber HE, Ingram JA, Davis DE, Hanley EN. Increased cell senescence is associated with decreased cell proliferation in vivo in the degenerating human annulus. Spine Journal [Internet]. 2009;9(3):210–5. Available from: http://dx.doi.org/10.1016/j.spinee.2008.01.012

14. Gruber HE, Ingram JA, Norton HJ, Hanley EN. Senescence in cells of the aging and degenerating intervertebral disc: Immunolocalization of senescence-associated β-galactosidase in human and sand rat discs. Spine (Phila Pa 1976). 2007;32(3):321–7.

15. Li P, Gan Y, Xu Y, Song L, Wang L, Ouyang B, et al. The inflammatory cytokine TNF-α promotes the premature senescence of rat nucleus pulposus cells via the PI3K/Akt signaling pathway. Sci Rep [Internet]. 2017 Mar 17 [cited 2020 Jul 13];7(1):42938–50. Available from: http://www.nature.com/scientificreports

16. Ashraf S, Santerre P, Kandel R. Induced senescence of healthy nucleus pulposus cells is mediated by paracrine signaling from TNF-α-activated cells. the FASEB journal [Internet]. 2021;35(9):e21795–810. Available from: http://www.ncbi.nlm.nih.gov/pubmed/34403508

17. Dimozi A, Mavrogonatou E, Sklirou A, Kletsas D. Oxidative stress inhibits the proliferation, induces premature senescence and promotes a catabolic phenotype in human nucleus pulposus intervertebral disc cells. Eur Cell Mater. 2015;30:89–103.

18. Gruber HE, Hoelscher GL, Ingram JA, Bethea S, Hanley EN. IGF-1 rescues human intervertebral annulus cells from in vitro stress-induced premature senescence. Growth Factors. 2008;26(4):220–5.

19. Forman HJ. Glutathione - From antioxidant to post-translational modifier. Arch Biochem Biophys. 2016 Apr 1;595:64–7.

20. Kim Y, Jang HH. Role of Cytosolic 2-Cys Prx1 and Prx2 in Redox Signaling. Antioxidants [Internet]. 2019 Jun 1 [cited 2022 Aug 16];8(6). Available from: /pmc/articles/PMC6616918/

21. Shields HJ, Traa A, van Raamsdonk JM. Beneficial and Detrimental Effects of Reactive Oxygen Species on Lifespan: A Comprehensive Review of Comparative and Experimental Studies. Front Cell Dev Biol [Internet]. 2021 Feb 11 [cited 2022 Jul 18];9:1–27. Available from: https://pubmed-ncbi-nlm-nih-gov.myaccess.library.utoronto.ca/33644065/

22. Liu Q, Tan Z, Xie C, Ling L, Hu H. Oxidative stress as a critical factor might involve in intervertebral disc degeneration via regulating NOXs/FOXOs. Journal of Orthopaedic Science. 2021 Nov 9;

23. Liu Y, Yu T, Ma XX, Xiang HF, Hu YG, Chen BH. Lentivirus-mediated TGF-β3, CTGF and TIMP1 gene transduction as a gene therapy for intervertebral disc degeneration in an in vivo rabbit model. Exp Ther Med. 2016;11(4):1399–404.

24. Hu B, Shi C, Xu C, Cao P, Tian Y, Zhang Y, et al. Heme oxygenase-1 attenuates IL-1β induced alteration of anabolic and catabolic activities in intervertebral disc degeneration. Sci Rep. 2016;6(415):1–14.

25. Wang D, Zheng H, Zhou W, Duan Z, Jiang S, Li B, et al. Mitochondrial Dysfunction in Oxidative Stress-Mediated Intervertebral Disc Degeneration. Orthop Surg [Internet]. 2022 Jun 8;(1665):1–14. Available from: https://onlinelibrary.wiley.com/doi/10.1111/os.13302

26. Ighodaro OM, Akinloye OA. First line defence antioxidants-superoxide dismutase (SOD), catalase (CAT) and glutathione peroxidase (GPX): Their fundamental role in the entire antioxidant defence grid. Alexandria Journal of Medicine [Internet]. 2018;54(4):287–93. Available from: https://doi.org/10.1016/j.ajme.2017.09.001

27. Zhu H, Chen G, Wang Y, Lin X, Zhou J, Wang Z, et al. Dimethyl fumarate protects nucleus pulposus cells from inflammation and oxidative stress and delays the intervertebral disc degeneration. Exp Ther Med [Internet]. 2020 Oct 27;20(6):1–10. Available from: http://www.spandidos-publications.com/10.3892/etm.2020.9399

28. Kudelko M, Chen P, Tam V, Zhang Y, Kong OY, Sharma R, et al. PRIMUS: Comprehensive proteomics of mouse intervertebral discs that inform novel biology and relevance to human disease modelling. Matrix Biol Plus [Internet]. 2021;12:100082. Available from: https://doi.org/10.1016/j.mbplus.2021.100082

29. Panebianco CJ, Dave A, Charytonowicz D, Sebra R, Iatridis JC. Single-cell RNA-sequencing atlas of bovine caudal intervertebral discs: Discovery of heterogeneous cell populations with distinct roles in homeostasis. FASEB Journal. 2021;35(11):1–21.

30. Hou G, Lu H, Chen M, Yao H, Zhao H. Oxidative stress participates in age-related changes in rat lumbar intervertebral discs. Arch Gerontol Geriatr [Internet]. 2014;59(3):665–9. Available from: http://dx.doi.org/10.1016/j.archger.2014.07.002

31. Feng C, Yang M, Lan M, Liu C, Zhang Y, Huang B, et al. ROS: Crucial Intermediators in the Pathogenesis of Intervertebral Disc Degeneration. Oxid Med Cell Longev [Internet]. 2017;2017:1–12. Available from: http://www.ncbi.nlm.nih.gov/pubmed/28392887

32. Upenieks A, Montgomery-Song A, Santerre JP, Kandel RA. Development of a Perfusion Reactor for Intervertebral Disk Regeneration. Tissue Eng Part C Methods [Internet]. 2022 Jan 1 [cited 2022 Jul 18];28(1):12–22. Available from: https://www-liebertpub-com.myaccess.library.utoronto.ca/doi/10.1089/ten.tec.2021.0216

33. Iu J, Santerre JP, Kandel RA. Inner and outer annulus fibrosus cells exhibit differentiated phenotypes and yield changes in extracellular matrix protein composition in vitro on a polycarbonate urethane scaffold. Tissue Eng Part A. 2014;20(23–24):3261–9.

34. Iu J, Santerre JP, Kandel RA. Towards engineering distinct multi-lamellated outer and inner annulus fibrosus tissues. Journal of Orthopaedic Research. 2018;36(5):1346–55.

35. Khan SY, Awad EM, Oszwald A, Mayr M, Yin X, Waltenberger B, et al. Premature senescence of endothelial cells upon chronic exposure to TNFα can be prevented by N-acetyl cysteine and plumericin. Sci Rep [Internet]. 2017 Jan 3 [cited 2022 Aug 21];7. Available from: /pmc/articles/PMC5206708/

36. Nisimoto Y, Diebold BA, Constentino-Gomes D, Lambeth JD. Nox4: A hydrogen peroxide-generating oxygen sensor. Biochemistry. 2014;53(31):5111–20.

37. Dou Y, Sun X, Ma X, Zhao X, Yang Q. Intervertebral Disk Degeneration: The Microenvironment and Tissue Engineering Strategies. Front Bioeng Biotechnol. 2021 Jul 20;0:490.

38. Chen C, Huang M, Han Z, Shao L, Xie Y, Wu J, et al. Quantitative T2 magnetic resonance imaging compared to morphological grading of the early cervical intervertebral disc degeneration: an evaluation approach in asymptomatic young adults. PLoS One [Internet]. 2014 Feb 3 [cited 2022 Aug 24];9(2). Available from: https://pubmed-ncbi-nlm-nih-gov.myaccess.library.utoronto.ca/24498384/

39. Suzuki S, Fujita N, Hosogane N, Watanabe K, Ishii K, Toyama Y, et al. Excessive reactive oxygen species are therapeutic targets for intervertebral disc degeneration. Arthritis Res Ther [Internet]. 2015;17(1):1–17. Available from: http://dx.doi.org/10.1186/s13075-015-0834-8

40. Feng C, Zhang Y, Yang M, Lan M, Liu H, Huang B, et al. Oxygen-Sensing Nox4 Generates Genotoxic ROS to Induce Premature Senescence of Nucleus Pulposus Cells through MAPK and NF-κB Pathways. Oxid Med Cell Longev [Internet]. 2017;2017:1–15. Available from: http://www.ncbi.nlm.nih.gov/pubmed/29147462

41. Guo Q, Zhu D, Wang Y, Miao Z, Chen Z, Lin Z, et al. Targeting STING attenuates ROS induced intervertebral disc degeneration. Osteoarthritis Cartilage. 2021;29(8):1213–24.

42. Richardson SM, Knowles R, Marples D, Hoyland JA, Mobasheri A. Aquaporin expression in the human intervertebral disc. J Mol Histol [Internet]. 2008 Jun 6 [cited 2022 Aug 17];39(3):303–9. Available from: https://link-springer-com.myaccess.library.utoronto.ca/article/10.1007/s10735-008-9166-1

43. Snuggs JW, Day RE, Bach FC, Conner MT, Bunning RAD, Tryfonidou MA, et al. Aquaporin expression in the human and canine intervertebral disc during maturation and degeneration. JOR Spine [Internet]. 2019 Mar 1 [cited 2022 Aug 17];2(1). Available from: /pmc/articles/PMC6686802/

44. Iwashita H, Castillo E, Messina MS, Swanson RA, Chang CJ. A tandem activity-based sensing and labeling strategy enables imaging of transcellular hydrogen peroxide signaling. Proc Natl Acad Sci U S A. 2021;118(9):1–9.

45. Kristiansen KA, Jensen PE, Møller IM, Schulz A. Monitoring reactive oxygen species formation and localisation in living cells by use of the fluorescent probe CM-H2DCFDA and confocal laser microscopy. Physiol Plant [Internet]. 2009 Aug 1 [cited 2022 Aug 17];136(4):369–83. Available from: https://onlinelibrary-wiley-com.myaccess.library.utoronto.ca/doi/full/10.1111/j.1399-3054.2009.01243.x

46. Møller IM, Jensen PE, Hansson A. Oxidative Modifications to Cellular Components in Plants. http://dx.doi.org.myaccess.library.utoronto.ca/101146/annurev.arplant58032806103946 [Internet]. 2007 May 1 [cited 2022 Aug 17];58:459–81. Available from: https://www-annualreviews-org.myaccess.library.utoronto.ca/doi/abs/10.1146/annurev.arplant.58.032806.103946

47. Krishnamoorthy L, Chang CJ. Exosomal NADPH Oxidase: Delivering Redox Signaling for Healing. Biochemistry [Internet]. 2018;57(27):3993–4. Available from: http://www.ncbi.nlm.nih.gov/pubmed/29889508

48. Bodega G, Alique M, Puebla L, Carracedo J, Ramírez RM. Microvesicles: ROS scavengers and ROS producers. J Extracell Vesicles [Internet]. 2019;8(1):1626654. Available from: https://doi.org/10.1080/20013078.2019.1626654

49. Johannessen W, Elliott DM. Effects of degeneration on the biphasic material properties of human nucleus pulposus in confined compression. Spine (Phila Pa 1976) [Internet]. 2005 Dec [cited 2022 Jul 18];30(24):E724–9. Available from: https://journals-lww-com.myaccess.library.utoronto.ca/spinejournal/Fulltext/2005/12150/Effects_of_Degeneration_on_the_Biphasic_Material.29.aspx

50. Zhao L, Tian B, Xu Q, Zhang C, Zhang L, Fang H. Extensive mechanical tension promotes annulus fibrosus cell senescence through suppressing cellular autophagy. Biosci Rep. 2019;39(4):1–10.

51. Meager A, Sampson LE, Grell M, Scheurich P. Development of resistance to tumour necrosis factor (tnfα) in kym-1 cells involves both tnf receptors. Cytokine. 1993;5(6):556–63.

52. Gregory AP, Dendrou CA, Attfield KE, Haghikia A, Xifara DK, Butter F, et al. TNF receptor 1 genetic risk mirrors outcome of anti-TNF therapy in multiple sclerosis. Nature 2012 488:7412 [Internet]. 2012 Jul 8 [cited 2022 Jul 18];488(7412):508–11. Available from: https://www.nature.com/articles/nature11307

53. Mutch DG, Powell CB, Kao MS, Collins JL. Resistance to cytolysis by tumor necrosis factor alpha in malignant gynecological cell lines is associated with the expression of protein(s) that prevent the activation of phospholipase A2 by tumor necrosis factor alpha. Cancer Res. 1992 Feb;52(4):866–72.

54. el Mahdani NE, Ameyar M, Cai Z, Colard O, Masliah J, Chouaib S. Resistance to TNF-Induced Cytotoxicity Correlates with an Abnormal Cleavage of Cytosolic Phospholipase A 2. The Journal of Immunology. 2000;165(12):6756–61.

55. Polla BS, Jacquier-Sarlin MR, Kantengwa S, Mariéthoz E, Hennet T, Russo-Marie F, et al. TNFα alters mitochondrial membrane potential in L929 but not in TNFα-resistant L929.12 cells: Relationship with the expression of stress proteins, annexin 1 and superoxide dismutase activity. Free Radic Res. 1996;25(2):125–31.

56. Dasgupta J, Subbaram S, Connor KM, Rodriguez AM, Tirosh O, Beckman JS, et al. Manganese superoxide dismutase protects from TNF-α-induced apoptosis by increasing the steady-state production of H2O2. Antioxid Redox Signal. 2006;8(7–8):1295–305.

57. Alkhatib B, Ban GI, Williams S, Serra R. IVD Development: Nucleus pulposus development and sclerotome specification. Curr Mol Biol Rep [Internet]. 2018 Sep;4(3):132–41. Available from: http://www.ncbi.nlm.nih.gov/pubmed/30505649

58. Du K, Yu Y, Zhang D, Luo W, Huang H, Chen J, et al. NFκB1 (p50) suppresses SOD2 expression by inhibiting FoxO3a transactivation in a mir190/phlpp1/akt-dependent axis. Mol Biol Cell. 2013;24(22):3577–83.

59. Jin LY, Lv ZD, Wang K, Qian L, Song XX, Li XF, et al. Estradiol Alleviates Intervertebral Disc Degeneration through Modulating the Antioxidant Enzymes and Inhibiting Autophagy in the Model of Menopause Rats. Oxid Med Cell Longev [Internet]. 2018;2018:7890291. Available from: http://www.ncbi.nlm.nih.gov/pubmed/30671175

60. Yang RZ, Xu WN, Zheng HL, Zheng XF, Li B, Jiang LS, et al. Involvement of oxidative stress-induced annulus fibrosus cell and nucleus pulposus cell ferroptosis in intervertebral disc degeneration pathogenesis. J Cell Physiol. 2021;236(4):2725–39.

61. Wang D, Zhang J, Sun Y, Lv N, Sun J. Long non-coding RNA NKILA weakens TNF-α-induced inflammation of MRC-5 cells by miR-21 up-regulation. Artif Cells Nanomed Biotechnol. 2020;48(1):498–505.

62. Liu B, Sun L, Liu Q, Gong C, Yao Y, Lv X, et al. A Cytoplasmic NF-κB Interacting Long Noncoding RNA Blocks IκB Phosphorylation and Suppresses Breast Cancer Metastasis. Cancer Cell [Internet]. 2015;27(3):370–81. Available from: http://dx.doi.org/10.1016/j.ccell.2015.02.004

63. He Y, Xiao Y, Yang X, Li Y, Wang B, Yao F, et al. SIRT6 inhibits TNF-α-induced inflammation of vascular adventitial fibroblasts through ROS and Akt signaling pathway. Exp Cell Res. 2017;357(1):88–97.

64. Zwacka RM, Stark L, Dunlop MG. NF-κB kinetics predetermine TNF-α sensitivity of colorectal cancer cells. Journal of Gene Medicine. 2000;2(5):334–43.

65. Dimri GP, Lee X, Basile G, Acosta M, Scott G, Roskelley C, et al. A biomarker that identifies senescent human cells in culture and in aging skin in vivo. Proc Natl Acad Sci U S A [Internet]. 1995 Sep 26;92(20):9363–7. Available from: http://www.ncbi.nlm.nih.gov/pubmed/7568133

66. Lee BY, Han JA, Im JS, Morrone A, Johung K, Goodwin EC, et al. Senescence-associated β-galactosidase is lysosomal β-galactosidase. Aging Cell. 2006;5(2):187–95.

67. Yang NC, Hu ML. The limitations and validities of senescence associated-β-galactosidase activity as an aging marker for human foreskin fibroblast Hs68 cells. Exp Gerontol. 2005;40(10):813–9.

68. Odgren PR, MacKay CA, Mason-Savas A, Yang M, Mailhot G, Birnbaum MJ. False-positive β-galactosidase staining in osteoclasts by endogenous enzyme: Studies in neonatal and month-old wild-type mice. Connect Tissue Res. 2006;47(4):229–34.

69. Kopp HG, Hooper AT, Shmelkov S v., Rafii S. β-galactosidase staining on bone marrow. The osteoclast pitfall. Histol Histopathol. 2007;22(7–9):971–6.

70. Piechota M, Sunderland P, Wysocka A, Nalberczak M, Sliwinska MA, Radwanska K, et al. Is senescence-associated β-galactosidase a marker of neuronal senescence? Oncotarget. 2016;7(49):81099–109.

71. Eble JA, de Rezende FF. Redox-relevant aspects of the extracellular matrix and its cellular contacts via integrins. Antioxid Redox Signal [Internet]. 2014 May 1 [cited 2022 Jul 15];20(13):1977–93. Available from: /pmc/articles/PMC3993061/

72. Xie J, Li B, Yao B, Zhang P, Wang L, Lu H, et al. Transforming growth factor-β1-regulated Fas/FasL pathway activation suppresses nucleus pulposus cell apoptosis in an inflammatory environment. Biosci Rep. 2020;40(2):1–8.

73. Chen HY, Ho YJ, Chou HC, Liao EC, Tsai YT, Wei YS, et al. TGF-β1 signaling protects retinal ganglion cells from oxidative stress via modulation of the HO-1/Nrf2 pathway. Chem Biol Interact. 2020 Nov 1;331:109249.

74. Lee YJ, Streuli CH. Extracellular Matrix Selectively Modulates the Response of Mammary Epithelial Cells to Different Soluble Signaling Ligands. Journal of Biological Chemistry. 1999 Aug 6;274(32):22401–8.

75. Lerche M, Elosegui-Artola A, Kechagia JZ, Guzmán C, Georgiadou M, Andreu I, et al. Integrin Binding Dynamics Modulate Ligand-Specific Mechanosensing in Mammary Gland Fibroblasts. iScience [Internet]. 2020 Mar 27 [cited 2022 Jul 15];23(3):100907–19. Available from: /pmc/articles/PMC7482011/

76. Zhang X, Wang X, Gao L, Yang B, Wang Y, Niu K, et al. TNF-а Induces Methylglyoxal Accumulation in Lumbar Herniated Disc of Patients With Radicular Pain. Front Behav Neurosci. 2021 Nov 23;15.

77. Hemanta D, Jiang X Xing, Feng Z Zhou, Chen Z Xian, Cao Y Wu. Etiology for Degenerative Disc Disease. Chinese Medical Sciences Journal. 2016 Sep 1;31(3):185–91.

78. Doraisamy R, Ramaswami K, Shanmugam J, Subramanian R, Sivashankaran B. Genetic risk factors for lumbar disc disease. Clin Anat [Internet]. 2021 Jan 1 [cited 2022 Jul 15];34(1):51–6. Available from: https://pubmed-ncbi-nlm-nih-gov.myaccess.library.utoronto.ca/32583875/

79. Cazzanelli P, Wuertz-Kozak K. MicroRNAs in Intervertebral Disc Degeneration, Apoptosis, Inflammation, and Mechanobiology. Int J Mol Sci [Internet]. 2020 May 20 [cited 2022 Jul 15];21(10):1–15. Available from: https://pubmed-ncbi-nlm-nih-gov.myaccess.library.utoronto.ca/32443722/

